# Myelination across cortical hierarchies and depths in humans and macaques

**DOI:** 10.1101/2025.02.06.636851

**Authors:** Monami Nishio, Xingyu Liu, Allyson P. Mackey, Michael J. Arcaro

**Affiliations:** Department of Psychology, School of Arts and Sciences, University of Pennsylvania, Philadelphia, Pennsylvania 19104, United States

**Keywords:** T1w/T2w, myelination, laminar architecture, cross-species

## Abstract

Myelination is fundamental to brain function, enabling rapid neural communication and supporting neuroplasticity throughout the lifespan. While hierarchical patterns of myelin maturation across the cortical surface are well-documented in humans, it remains unclear which features reflect evolutionarily conserved developmental processes versus human-characteristic adaptations. Moreover, the laminar development of myelin across the primate cortical surface, which shapes hierarchies and supports functions ranging from sensory integration to network communication, has been largely unexplored. Using neuroimaging to measure the T1-weighted/T2-weighted ratio in tissue contrast as a proxy for myelin content, we systematically compared depth-dependent trajectories of myelination across the cortical surface in humans and macaques. We identified a conserved “inside-out” pattern, with deeper layers exhibiting steeper increases in myelination and earlier plateaus than superficial layers. This depth-dependent organization followed a hierarchical gradient across the cortical surface, progressing from early maturation in sensorimotor regions to prolonged development in association areas. Humans exhibited a markedly extended timeline of myelination across both cortical regions and depths compared to macaques, allowing for prolonged postnatal plasticity across the entire cortical hierarchy — from sensory and motor processing to higher-order association networks. This extended potential for plasticity may facilitate the shaping of cortical circuits through postnatal experience in ways that support human-characteristic perceptual and cognitive capabilities.

## Introduction

Myelination is integral to brain function, enhancing neural communication, supporting synaptic plasticity, and fostering functional specialization^1^. By increasing axonal conduction velocity and metabolic efficiency, myelin enables rapid and reliable information transfer necessary for sensory processing, motor control, and higher-order cognition^2–4^. Particularly in larger-brained, gyrencephalic great apes, extensive myelination shapes complex neural circuits underlying advanced perceptual and cognitive capabilities throughout postnatal development^1^.

The maturation of myelin in humans follows a hierarchical trajectory that parallels the functional organization of the adult cortex^5,6^. Primary sensory and motor regions, associated with rapid, reliable processing, are myelinated early, while association cortices, which support integrative processing across distributed networks, show protracted development^7^. Human neuroimaging studies, with their capacity for whole-brain coverage, have been pivotal in elucidating this hierarchical architecture, revealing broad patterns of myelination across distributed cortical networks^7^. However, the evolutionary origins of these patterns remain unresolved. Evidence from nonhuman primates suggests similar regional patterns of myelination^6^, but these studies typically rely on *ex vivo* methods with sparse sampling across the cortical surface, making it difficult to capture global hierarchical relationships. Consequently, it remains unclear which aspects of these myelination patterns reflect developmental processes conserved across primates and which might represent adaptations unique to humans.

The laminar organization of cortex, with its systematic variation in cell type composition and connectivity patterns, is a defining feature of cortical hierarchies^8,9^. This structure underpins processes ranging from sensory integration to higher-order cognition^8,9^. Despite its crucial role in organizing hierarchical networks, the developmental trajectory of laminar myelination and how it shapes functioning across the cortical hierarchy remain poorly understood. Recent studies in rodents suggest that myelination follows an “inside-out” pattern, where deeper cortical layers mature earlier than superficial layers^10–13^. For example, immunohistochemical studies of myelin basic protein (MBP) in mice show that the first myelinated axons appear in layers V–VI^10^. Whether this developmental pattern represents a conserved feature across mammals remains an open question, particularly for primates, which exhibit an expanded cortical hierarchy^14^ and unique developmental features of the cortical plate^15^. In primates, especially humans, the disproportionate expansion of superficial layer neurons^15^ and their increased connectivity complexity^16^ contribute to a laminar architecture that is both highly differentiated and specialized^17^. Understanding how an ‘inside-out’ pattern of myelination is elaborated to support these unique developmental characteristics of the primate brain remains a critical area of investigation. Recent human imaging studies align with an “inside-out” pattern of myelo-development, suggesting that deeper cortical layers exhibit a steeper rate of myelination increase compared to superficial layers during adolescence^18,19^. However, the age range tested was too restricted to determine whether this steeper increase during adolescence represents simultaneous maturation at different rates or a hierarchical variation in the timing of plateauing across layers. Adding to this complexity, regional variability in the laminar maturation of cytoarchitecture has been observed within sensory cortices of marmoset^20^, leaving open the question of whether the temporal pattern of laminar myelination similarly varies across cortical regions in primates.

While patterns of myelination across species provide a foundation for understanding conserved mechanisms of cortical development, humans exhibit exceptionally prolonged maturation of cortical myelination compared to other primates^6,21^. In macaques, myelination largely stabilizes within the first three years of life^21,22^, while our closest living relatives, chimpanzees, exhibit a more extended timeline, with myelination continuing to increase until reaching adult-like levels at sexual maturity (around ages 4-6 years)^6^. In humans, however, myelination progresses along a markedly prolonged trajectory, continuing into the third decade and beyond^6,23^. This extended developmental timeline is thought to enable greater plasticity, providing a longer window for environmental influences to shape cortical circuits during postnatal development^5,6^. The uniquely prolonged cortical development in humans suggests possible species-specific modifications to depth-dependent myelination patterns. Superficial layers, with their dense cortico-cortical connections, may undergo especially prolonged development, reflecting their central roles in integrating information across distributed networks^9,17^. While much attention has been directed toward the evolutionary expansion of superficial layers, deeper layers may also exhibit prolonged developmental timelines, as their structural maturation supports thalamocortical and subcortical pathways essential for sensory processing, attention, and other cognitive functions^9^. Resolving the timecourse of these developmental patterns requires systematic cross-species comparisons to determine how conserved developmental frameworks interact with human-characteristic adaptations.

Prior rodent and monkey studies have offered important insights into myelination but were limited by narrow age ranges and restricted cortical sampling, hindering a comprehensive understanding of how myelination progresses across cortical regions and depths throughout development. Advances in neuroimaging, particularly the use of the T1-weighted/T2-weighted (T1w/T2w) ratio as a proxy for myelin content, enable non-invasive, whole-brain examination of cortical myelination across broad age ranges in multiple primate species^24–26^.

Building on this foundation, the current study systematically examines region-wise and depth-dependent trajectories of myelin development across the cortical surface in humans and macaques. We developed a cross-primate analysis pipeline to assess T1w/T2w ratio patterns across cortical regions and depths, validated with histological data from macaques. This approach allows us to identify both shared and species-specific patterns of myelin development, providing insights into the evolutionary adaptations that shape primate brain organization. By comparing regional and depth-dependent myelination trajectories across macaques and humans, we aim to elucidate how conserved developmental mechanisms interact with human-characteristic adaptations, shedding light on the neural correlates of advanced perceptual and cognitive functions.

## Results

### Regional and depth-dependent T1w/T2w ratio variability in macaques and humans

To investigate the spatial distribution of cortical myelination and establish a reference for later-stage myelination patterns, we first analyzed T1w/T2w ratios across cortical regions and depths in late adolescent macaques (Fig. 1A, ages 2-3 years, *N* = 25) and young adult humans (Fig. 1B, ages 22-36 years, *N* = 1,113).

**Figure 1.**
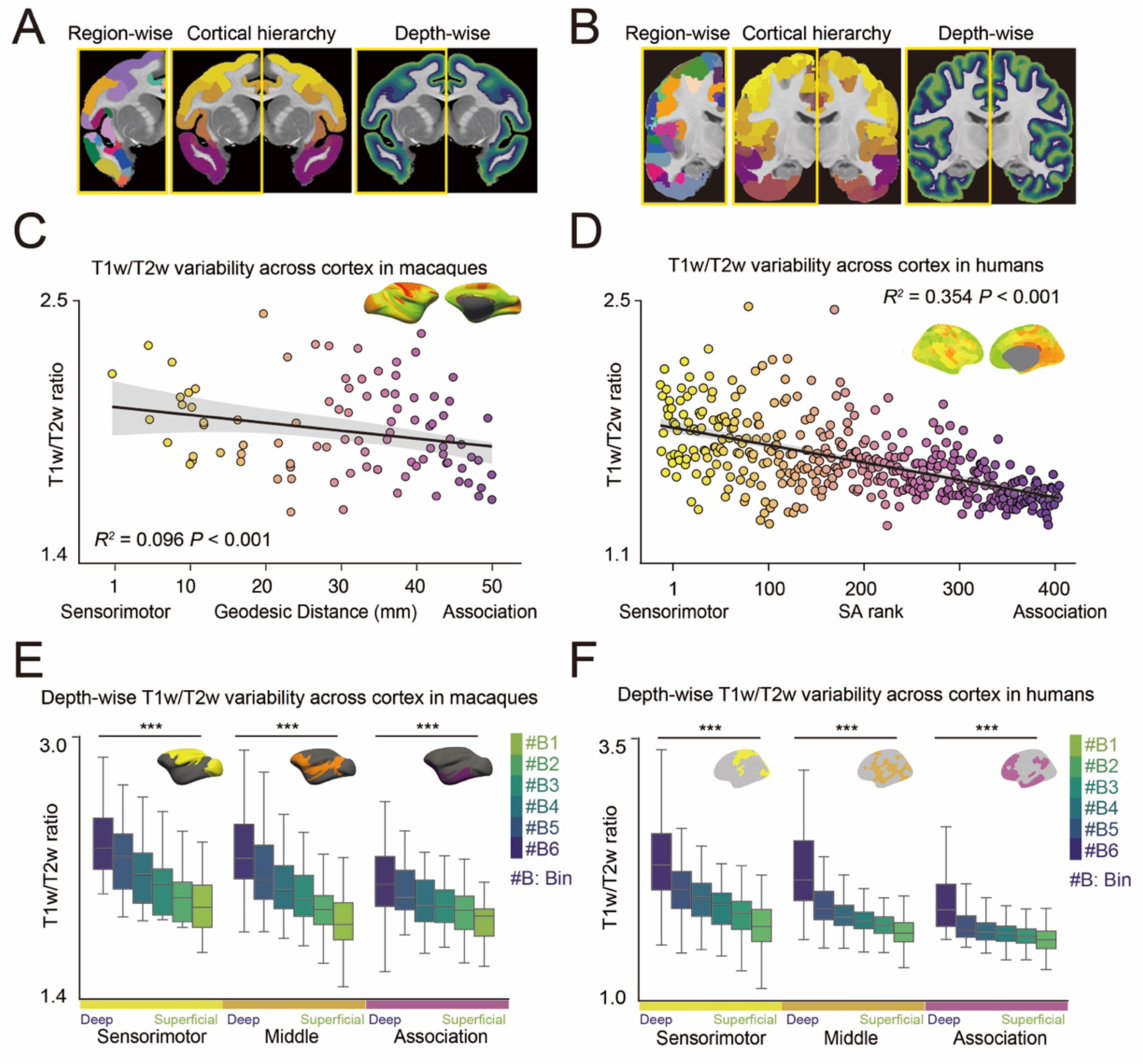
T1w/T2w ratio across cortical hierarchy and depth in macaques and humans. **(A, B)** Schematic of the analytical approach for evaluating the T1w/T2w ratio across cortical regions and depths in macaques (A) and humans (B). The left panel displays the discrete parcels of CHARM level 6^27,28^ for macaques and Schaefer 400^29^ for humans. The middle panel labels the parcels based on geodesic distance for macaques or the sensorimotor-association axis for humans, with colors transitioning from yellow (sensorimotor) to purple (association). The right panel visualizes the laminar organization, with colors transitioning from deep blue (deep layers) to light green (superficial layers). **(C, D)** Distribution of T1w/T2w ratios along the geodesic distance for macaques (C, *R^2^*= 0.096, *P* < 0.001) and along the sensorimotor-association (S-A) axis for humans (D, *R^2^* = 0.354, *P* < 0.001). **(E, F)** Cortical depth-wise T1w/T2w ratios within sensorimotor, middle, and association regions in macaques (E) and humans (F); ANOVA \*\*\**P* < 0.001.

To establish a common framework for comparing cortical development between macaques and humans, we parcellated the cortex using species-specific anatomical templates: the CHARM level 6 template^27,28^, containing 139 cortical parcels, for macaques and the Schaefer 400 template^29^ for humans. Cortical regions were then grouped along the sensorimotor-association (S-A) axis, a primary organizational gradient spanning from sensory and motor cortices to transmodal, heteromodal, and paralimbic association cortices^30^. While this axis is well-characterized in humans, it remains less validated in macaques. Recent work has described a similar axis in macaques using tracer connectome data^31^, but this approach is limited by incomplete cortical coverage, particularly for association regions, making it insufficient for whole-brain analysis. To address this limitation, we employed an alternative measure of the S-A axis in macaques, based on geodesic distance along the cortical surface from predefined association areas^31^. This measure showed strong correspondence with both the tracer-derived S-A axis in macaques (Fig. S1A-C) and the multimodal-derived S-A axis in humans (Fig. S1D-F)^42^, establishing a framework for direct comparisons of the entire cortical hierarchy between species. To assess broad regional patterns, we categorized the cortical parcels into sensorimotor, middle, and association regions based on geodesic distance from association regions^31^ in macaques and the multimodal-derived S-A axis^30^ for humans. In both macaques and humans, the T1w/T2w ratios were higher in sensorimotor regions than in association regions, showing a linear decrease along the S-A axis (Fig. 1C; Macaques *R^2^* = 0.096, *P* < 0.001, Fig. 1D; Humans *R^2^*= 0.354, *P* < 0.001), replicating past work^32^. These results illustrate a conserved gradient in myelination, with sensorimotor regions exhibiting higher myelination than association cortices in both primate species.

To investigate depth-dependent variations in myelination across this cortical hierarchy, we segmented the gray matter into six equivolumetric bins using LayNii^33^. This approach was necessary because the resolution of anatomical MRI does not allow precise delineation of histological cortical layers. By applying this binning approach, we established a consistent method for analyzing depth-dependent myelination patterns across cortical regions and species. In both macaques and humans, deeper cortical bins exhibited significantly higher T1w/T2w ratios than superficial bins throughout the cortical hierarchy, spanning from sensorimotor to association regions (Fig. 1E; Macaques, 1F; Humans, all ANOVAs *F* > 9.236, *P* < 0.001), replicating prior findings^18,34,35^. To validate the T1w/T2w ratio as a proxy for depth-specific myelin content, we examined layer-specific expression of myelin basic protein (MBP) in macaques. We found a significant correlation between MBP expression and the T1w/T2w ratio across both the cortical surface and depth (see Supplementary Results, Fig. S2). Together, these results provide a robust framework for cross species comparisons of myelination and establish a foundation for studying its maturation across cortical regions and depths.

### Hierarchical development of T1w/T2w ratio along the sensorimotor-association axis is protracted in humans compared to macaques

Having characterized these myelination patterns, we next examined their region-wise developmental trajectories by analyzing the T1w/T2w ratio in macaques from 1 month to 3 years (*N* = 34), and in humans from 5.5 to 36 years (*N* = 1,730) (Fig. 2A). These age ranges contained overlapping developmental stages, as macaque brain development occurs approximately four times faster than human development based on a broad spectrum of established neurodevelopmental metrics^36^. We hypothesized that the developmental trajectories of myelination in both species would recapitulate the hierarchical organization of adult cortex, with earlier maturation in sensorimotor regions compared to association regions. However, we expected humans to exhibit more protracted developmental trajectories. We quantified developmental changes using Generalized Additive Models (GAM)^37^, calculating the differential in *R^2^* values between the full and reduced models excluding age effects. The direction of developmental change (*R*^2^_partial_) was determined by the sign of the average first derivative of the smooth age function.

**Figure 2.**
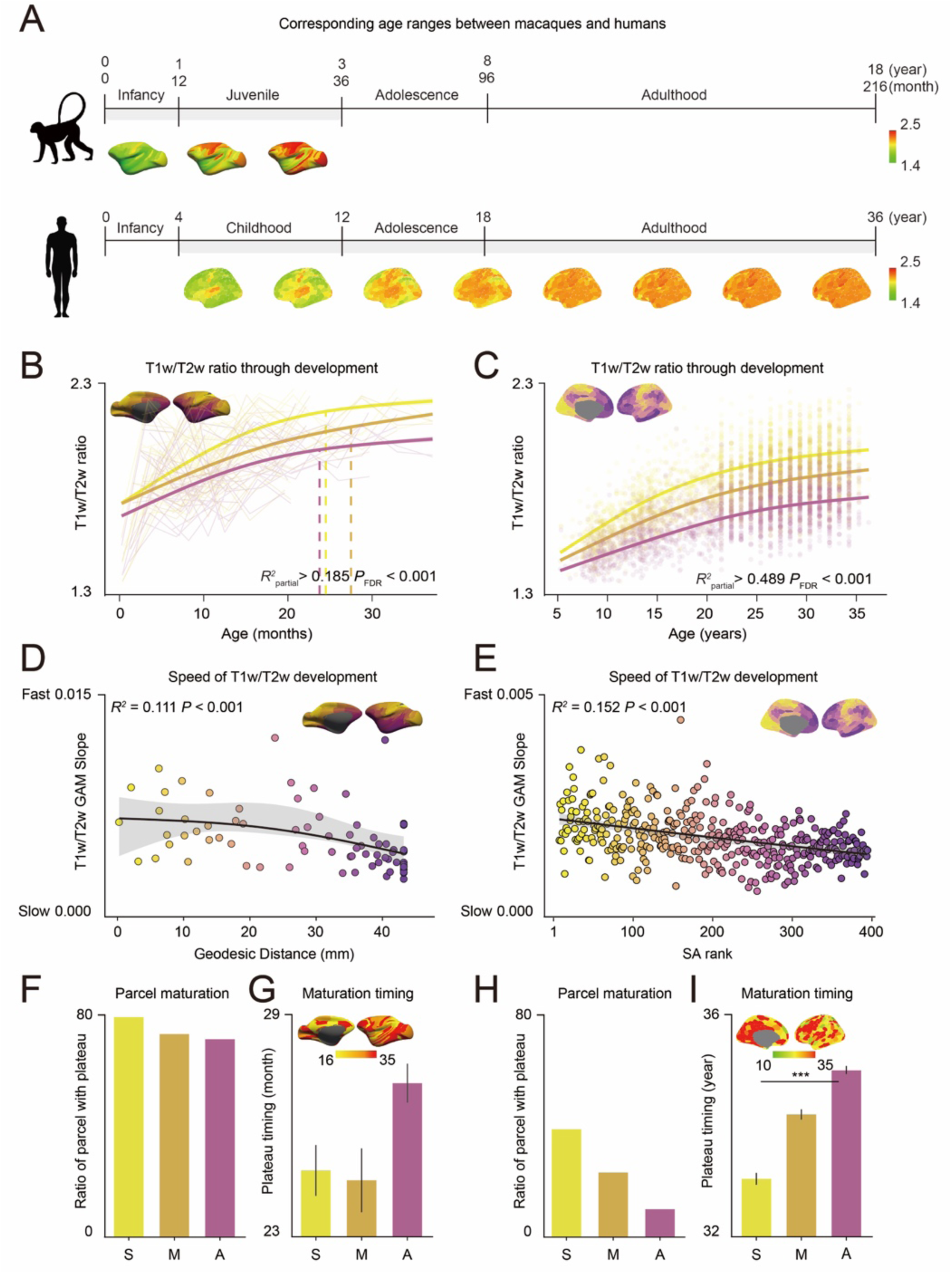
Development of T1w/T2w ratio across cortical hierarchy in macaques and humans. **(A)** Age ranges for the macaques and humans overlapped based on the 4:1 age scaling ratio. **(B, C)** Developmental trajectories of the T1w/T2w ratio for sensorimotor (yellow), middle (orange), and association (purple) cortical areas for macaques (B) and humans (C). Solid lines illustrate Generalized Additive Model (GAM)-predicted fits along with their 95% confidence intervals (B; Sensorimotor; *R*^2^_partial_ = 0.317, *P_FDR_* < 0.001, Middle; *R^2^* _partial_ = 0.195, *P_FDR_* < 0.001, Association; *R^2^*_partial_ = 0.185, *P_FDR_* < 0.001, C; Sensorimotor; *R^2^*_partial_ = 0.498, *P_FDR_* < 0.001, Middle; *R^2^*_partial_ = 0.539, *P_FDR_* < 0.001, Association; *R^2^* _partial_ = 0.489, *P_FDR_* < 0.001). Dashed lines indicate the age at which slope of the T1w/T2w ratio versus age becomes non-significant. **(D, E)** GAM slopes along the geodesic distance for macaques (D, *R^2^* = 0.111, *P* < 0.001) and along the sensorimotor-association (S-A) axis for humans (E, *R^2^* = 0.152, *P* < 0.001). The GAM-predicted fit is displayed with a 95% confidence interval (D; *R²* = 0.111, *P* < 0.001, E; *R² = 0.152, P < 0.001*). **(F, H)** Proportion of regions reaching plateau for sensorimotor (S), middle (M) and association (A) cortex in macaques (F) and humans (H). **(G, I)** Average timing of plateau for cortical regions in sensorimotor (S), middle (M) and association (A) regions for macaques (G; in months, ANOVA *F* = 1.353, *P* = 0.262) and humans (I; in years, ANOVA *F* = 14.725, *P* < 0.001). ANOVA \*\*\**P* < 0.001.

In macaques, the T1w/T2w ratio increased across development for all cortical regions, including sensorimotor, middle, and association regions, before plateauing around 24-30 months, corresponding developmentally to approximately 8-10 years old in humans based on broader metrics of brain maturation^36^ (Fig. 2B, all *R*^2^_partial_ > 0.185, *P_FDR_* < 0.001). In contrast, the T1w/T2w ratio in humans continued to increase with no detected plateau in any cortical region, even in individuals up to 36 years old, a developmental period far exceeding the corresponding age range analyzed in macaques (Fig. 2C, all *R^2^*_partial_ > 0.489, *P_FDR_* < 0.001). These results highlight the markedly prolonged developmental trajectory of myelination in humans compared to macaques.

The rate of myelination development varied across cortical regions, reflecting the hierarchical organization of the cortex in both macaques and humans. In both species, steeper T1w/T2w ratio increases were observed in sensorimotor regions compared to association regions, with sensorimotor regions plateauing earlier along this gradient (Fig. 2D, *R*² = 0.111, *P* < 0.001; Fig. 2E, *R*² = 0.152, *P* < 0.001). This developmental hierarchy was further supported by characterizing the timing and extent of plateaus across the S-A axis. In both macaques (Fig. 2F) and humans (Fig. 2H), a greater proportion of parcels in sensorimotor regions reached a plateau compared to association regions. The plateau occurred at an earlier age in sensorimotor and middle regions compared to association regions for both species, though this timing difference was statistically significant only in humans (macaques: Fig. 2G, ANOVA *F* = 1.353, *P* = 0.262; humans: Fig. 2I, ANOVA *F* = 14.725*, P* < 0.001). Supporting the robustness of these findings in humans, the hierarchical progression of T1w/T2w ratio development from sensorimotor to association regions persisted when the S-A axis was defined using geodesic distance from the default mode network, as done in the macaques’ analysis (Fig. S3).

Species differences in myelination trajectories were most evident in quantifying the proportion of regions reaching a plateau. In macaques, over 70% of cortical regions plateaued by age 3, corresponding to around 12 years in humans (Fig. 2F). In contrast, by age 36, fewer than 50% of human sensorimotor cortical regions, and only 20% of association regions had plateaued (Fig. 2H). These findings highlight the extended development of myelination in humans compared to macaques, beyond even the 4:1 scaling of broader brain maturation metrics^36^. This disproportionate extended development is particularly evident in association regions, emphasizing the protracted maturation of higher-order cortical areas in humans.

### Deeper layers plateau earlier than superficial layers in macaques and humans

We next examined the development of T1w/T2w ratio across cortical depths to understand laminar patterns of myelination and their relationship to cortical hierarchy in macaques and humans (Fig. 3A; Macaques, 3B; Humans). In both species, deeper layer bins exhibited significantly steeper developmental slopes than superficial layer bins across all cortical regions (Fig. 3C; Macaques all ANOVAs *F* > 7.734, *P* < 0.001, 3E; Humans all ANOVAs *F* > 20.327, *P* < 0.001). The magnitude of this cortical depth-dependent development varied by cortical region, with larger differences between superficial and deep bins in sensorimotor regions compared to association regions (Fig. 3D; Macaques *R²* = 0.061, *P* = 0.014, 3F; Humans *R²* = 0.120, *P* < 0.001).

**Figure 3.**
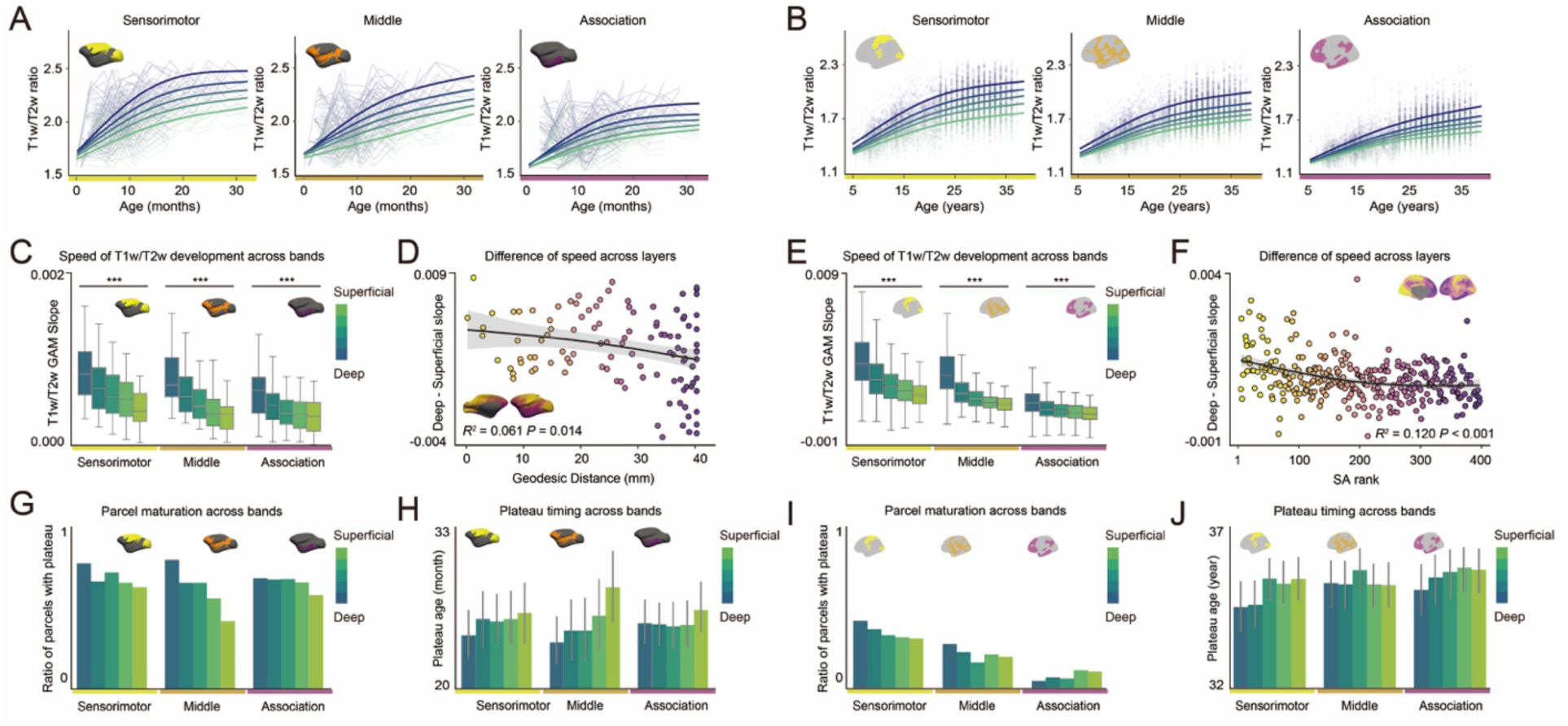
T1w/T2w ratio across cortical depth in macaques and humans. **(A, B)** T1w/T2w ratios and Generalized Additive Model (GAM)-predicted fits for cortical depth bin in sensorimotor, middle and association regions for macaques (A) and humans (B). **(C, E)** Mean GAM developmental slopes for each cortical depth bin for sensorimotor, middle, and association cortical regions for macaques (C; Sensorimotor; *F* = 11.295, *P* < 0.001, Middle; *F* = 11.947, *P* < 0.001, Association; *F* = 7.734, *P* < 0.001) and humans (D; Sensorimotor; *F* = 20.327, *P* < 0.001, Middle; *F* = 55.301, *P* < 0.001, Association; *F* = 99.100, *P* < 0.001). \*\*\**P* < 0.001. **(D, F)** Slope differences between the deepest and the most superficial bins along the geodesic distance in macaques (D) and along the sensorimotor-association (S-A) axis in humans (F), with GAM-predicted fits and their 95% confidence intervals (D; *R²* = 0.061, *P* = 0.014, F; *R² =* 0.120*, P* < 0.001). **(G, I)** Proportion of cortical parcels with plateau at each cortical depth bin for sensorimotor, middle, and association regions for macaques (G) and humans (I). **(H, J)** Average timing of plateau in each cortical depth bin for sensorimotor, middle, and association regions for macaques (H; Sensorimotor; *F* = 0.355, *P* = 0.840, Middle; *F* = 1.627, *P* = 0.170, Association; *F* = 0.315, *P* = 0.868) and humans (J; Sensorimotor; *F* = 1.666, *P* = 0.156, Middle; *F* = 0.971, *P* = 0.423, Association; *F* = 0.892, *P* = 0.468).

The depth-dependent developmental pattern was further supported by analyses of the plateau timing. In both species, a greater proportion of cortical regions in deeper bins reached a plateau compared to superficial bins (Fig. 3G; Macaques, 3I; Humans). These plateaus tended to occur earlier in deeper compared to superficial bins, though the timing difference was not significant (Fig. 3H; Macaques, ANOVAs *F* < 1.627, *P* > 0.170, 3J; Humans ANOVAs *F* < 1.666, *P* > 0.156). Together, these findings reveal a conserved “inside-out” pattern of myelination across cortical depths, highlighting depth-dependent developmental dynamics along the cortical sensorimotor-association axis and across primate species.

Species differences in myelination trajectories were evident across all cortical layers. In macaques, over 40% of cortical regions reached a plateau by age 3, even in the most superficial layers. In contrast, in humans, fewer than 45% of regions reached a plateau by age 36, even in the deepest layers. These findings suggest that, compared to macaques, humans maintain a prolonged window of myelination plasticity across cortical layers, not just superficial.

## Discussion

This study provides a comprehensive comparison of cortical myelination development in humans and macaques, highlighting both shared and species-specific patterns in the myelination trajectories across cortical regions and depths. Our analysis revealed a conserved hierarchical pattern of myelination across species, progressing from primary sensorimotor regions to association cortical areas. In parallel, we identified an “inside-out” pattern of laminar development, with deeper cortical layers exhibiting a steeper increase in myelination and reaching their plateaus earlier than superficial layers. Notably, these developmental patterns are prolonged in humans compared to closely related non-human primates, both across cortical regions and depths. This extended timeline in humans reflects species-specific adaptations that elaborate on conserved principles of cortical organization, providing an expanded window for postnatal development and experiential learning.

Comparative studies using histology and RNA, sampling from select cortical areas, have consistently demonstrated that non-human primates, including macaques^22^ and chimpanzees^6^, reach a myelination plateau earlier than humans. Using neuroimaging to map myeloarchitectural development across the entire cortical surface, our findings confirmed these observations, showing that the T1w/T2w ratio of human cortical areas across the sensorimotor-association axis continues to increase well into adulthood, long after the corresponding developmental period in non-human primates (Fig. 4B; sensorimotor regions, 4C; association regions). Notably, we found that T1w/T2w ratio development is prolonged in both deep and superficial layers (Fig. 4D; deep layers, 4E; superficial layers), suggesting unique adaptations in humans for both cortico-cortical connectivity in superficial layers and thalamocortical and subcortical connectivity in deeper layers. The prolonged period of myelination in humans likely supports an extended window for experience-dependent plasticity, potentially supporting the development of complex cognitive abilities and adaptability throughout life^38^. This finding aligns with the neuroconstructivist framework, which posits that the evolution of neocortex primarily reflects enhanced capacity for learning and plasticity rather than the maximization of pre-specified core knowledge (i.e., hyper-specialization)^39^.

**Figure 4.**
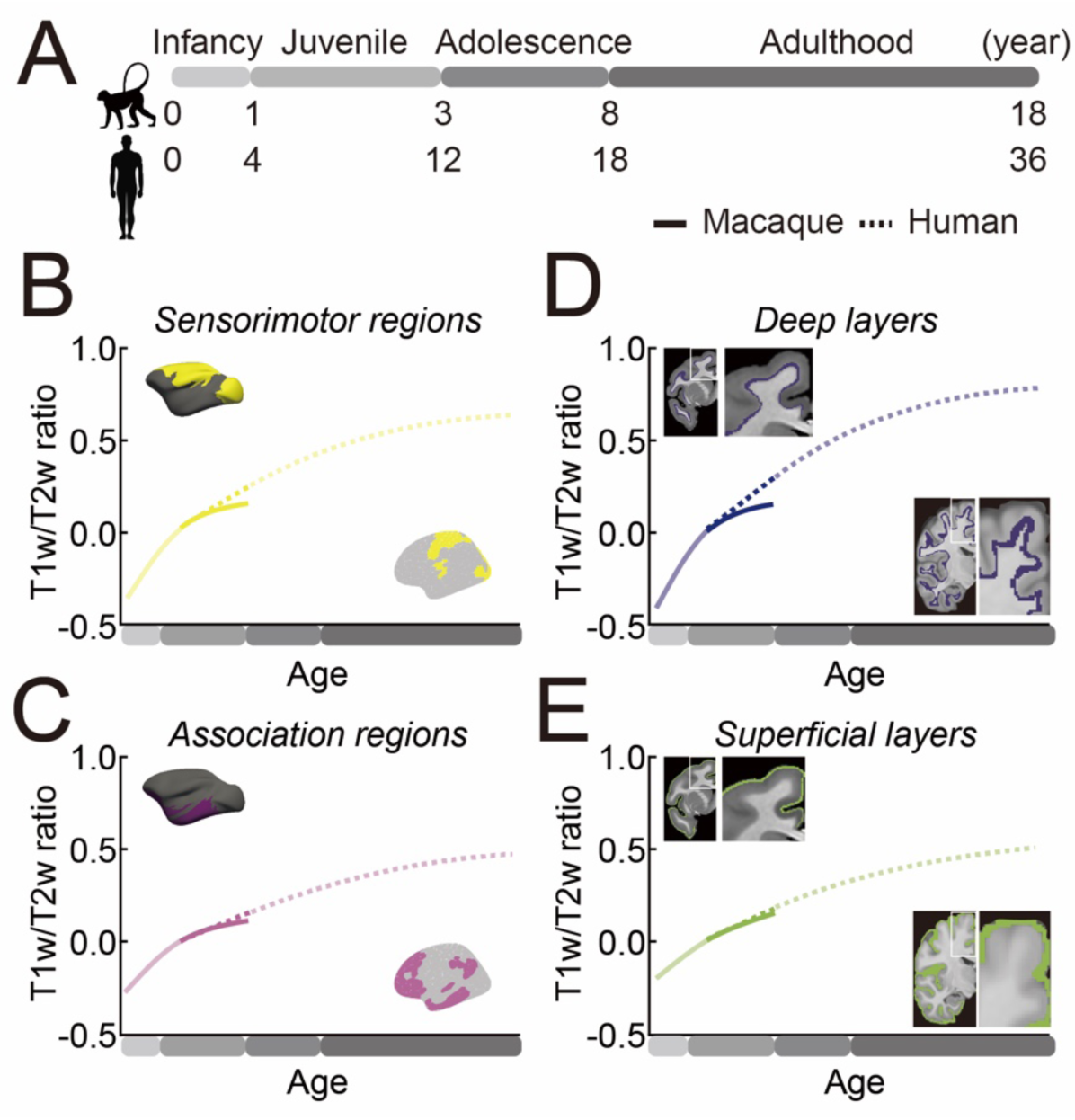
Summary of regional and cortical depth T1w/T2w Ratio Development in Humans and Macaques. **(A)** Age correspondence between macaques and humans assuming a 4:1 scaling. **(B, C)** T1w/T2w ratio developmental trajectories across sensorimotor (B, yellow) and association (C, purple) regions for macaques (straight) and humans (dotted). **(D, E)** T1w/T2w developmental trajectories across deep (D, dark blue) and superficial (E, green) cortical layers for macaques (straight) and humans (dotted). T1w/T2w values are adjusted relative to 1 year old (macaque) and 4 years old (human) as the baseline (y = 0). The age spans that overlap between macaques and humans are shown in solid colors, while the non-overlapping age span are plotted in diluted colors.

The extended development observed in humans is anchored to conserved processes shared with macaques. Both species display a hierarchical progression of myelination development across cortex, advancing from sensorimotor to association regions. These findings converge with prior research demonstrating a correspondence between functional connectivity and intracortical myelin in humans, suggesting that myelination patterns may reflect regional functional specialization^40^. Within this evolutionary framework, the human trajectory, while remarkable in its degree of protracted myelination, is not extraordinary, as it builds on conserved mechanisms underlying cortical development across primates^41^. This protracted postnatal maturation of association cortices in primates provides a scalable framework for supporting species-specific adaptation, such as the development of complex perceptual and cognitive functions in humans.

Previous studies indicate that deeper cortical layers exhibit a steeper rate of increase in myelination compared to superficial layers during adolescence^18,19^. However, mapping the entire trajectory of myelination in humans has proven challenging, as myelination maturation spans several decades. By analyzing macaque data, where myelination plateaus are observable within a shorter developmental window, we found that deeper layers not only exhibit a steeper increase in myelination but also reach a plateau earlier than superficial layers. This pattern was also observed in humans, although fewer than half of their cortical regions reached a plateau by age 36. These findings suggest that the “inside-out” pattern of cortical myelination is conserved across primates, with adaptations in humans. The functional implications of this pattern vary across cortical depths. Earlier maturation of deeper layers, which contain extensive subcortical and corticothalamic projection neurons, likely supports essential sensory processing and motor control during early postnatal development. In contrast, the prolonged development of superficial layers, predominantly containing cortico-cortical connections, extends the window for postnatal experiential learning and adaptation. This extended timeline may facilitate the refinement of integrative functions critical for higher-order perception and cognition, enabling the advanced cognitive flexibility and complexity characteristic of human behaviors.

Our study relied on measurements of T1w/T2w ratio using neuroimaging. While the T1w/T2w ratio is widely used as an indirect proxy for myelin content, limitations have also been noted^42–44^. In particular, its validity as a depth-wise myelin marker has not yet been fully established. Using a newly available dataset of layer-specific MBP RNA expression in macaques^45^, we demonstrated a clear alignment between MBP expression and T1w/T2w ratio across cortical depth (see Supplementary Results). This finding supports the use of the T1w/T2w ratio as a reliable depth-specific myelin marker across primates.

By revealing both common and human-characteristic features of myelination, our findings contribute to a broader understanding of human brain development and evolution. These results emphasize the value of comparative studies in elucidating both conserved principles and species-specific adaptations in brain development. The prolonged period of myelination in humans, particularly in superficial layers linked to cortico-cortical connectivity, may provide an extended window for environmental influences to shape higher-order association cortices crucial for human-characteristic behaviors such as abstract reasoning and social cognition. However, our findings demonstrate that this extended period of myelination in humans is not confined to these higher-order regions and associated functions. Instead, the protracted maturation of myelination extends beyond what would be predicted by the established four-fold difference in developmental timing between macaques and humans^36^, and encompasses the entire cortical surface and depths, including regions and layers critical for both sensory and integrative processes. This comprehensive extension reflects human-characteristic adaptations across the cortical hierarchy, supporting the advanced perceptual and cognitive capabilities characteristic of human behaviors.

## Acknowledgements

This work was supported by a Whitehall Foundation grant (to M.J.A.), NIH grants P30EY12196, R01EY025670, R01NS123778 (to M.J.A.), and NSF grants 2045095 (to A.P.M.). M.N. was supported by Quad Fellowship and Nakajima Foundation Scholarship. The funders had no role in the study design, data collection and interpretation, or the decision to submit the work for publication.

## Author Contributions

M.N., X.L. and M.J.A. conceptualized the study; M.N. and X.L. analyzed the data and prepared the figures; M.N., X.L., A.P.M. and M.J.A. interpreted the results; M.N., X.L. and M.J.A. wrote the manuscript; A.P.M. and M.J.A. supervised M.N; all authors provided feedback on the manuscript.

## Declaration of interest

The authors declare no competing interests.

## Declaration of generative AI and AI-assisted technologies

During the preparation of this work, the author used ChatGPT 3.5 to draft the first version of the manuscript. After using this tool, all authors edited the content substantially and take full responsibility for the content of the publication.

## STAR★Methods

### Resource Availability

#### Lead contact

Further information and requests for resources should be directed to and will be fulfilled by the lead contact, Monami Nishio.

#### Materials Availability

N/A

#### Data and Code Availability

- All data reported in this study will be publicly available as of the date of publication.
- All original code will be publicly available as of the date of publication.
- Any additional information required to reanalyze the data reported in this paper is available from the lead contact upon request.

### Participants

#### Humans

We analyzed neuroimaging data from 1,730 healthy participants: 617 aged 5.58–21.92 years (54.8% female) from the Human Connectome Project in Development (HCP-D; Release 2.0)^7^, and 1,113 participants aged 22-36 years (54.4% female) from the Human Connectome Project (HCP; S1200 release)^46^. We excluded 13 HCP-D participants due to missing T1w scans or anatomical anomalies, resulting in a final sample of 617 youth.

#### Macaques

We analyzed neuroimaging data from 34 rhesus monkeys (Macaca mulatta, 41.2% female) from the UNC-Wisconsin Rhesus Macaque Neurodevelopment Database^47^. Monkeys were reared and housed at the Harlow Primate Laboratory (HPL) at the University of Wisconsin-Madison. Each monkey underwent longitudinal scanning (five timepoints, except one monkey with four), with intervals based on age at first scan.

For single-nucleus RNA sequencing, we analyzed data from two monkeys as published in Chen et al., 2023^45^. Left hemispheres were collected from two male cynomolgus monkeys (*M. fascicularis*; #1, 6-year-old, 4.2 kg; #2, 4-year-old, 3.7 kg).

### Image Acquisition

#### Humans

High-resolution T1w MRI images were acquired on a 3T Siemens Prisma with a 32 channel head coil using a 3D multiecho MPRAGE sequence^48,49^ (0.8-mm isotropic voxels, TR/TI = 2500/1000 ms, TE = 1.8/3.6/5.4/7.2 ms, flip angle = 8°, in-plane (iPAT) acceleration factor of 2, TA = 8:22, up to 30 reacquired TRs). Structural T2w images were acquired using the variable-flip-angle turbo-spin-echo 3D SPACE sequence^50^ (0.8 mm isotropic voxels, TR/TE = 3200/564 ms; same in-plane acceleration, TA = 6:35, up to 25 reacquired TRs). Both pre scan normalized and non-normalized reconstructions were generated, with non-normalized versions used for subsequent processing^51^ and both used to estimate the B1– receive field for motion correction between T1w and T2w images. Only the first two T1w echoes were used due to artifacts in later echoes^52^. Volumetric navigators (vNavs) were embedded in T1w and T2w sequences for prospective motion correction and selective k-space reacquisition^53^, reducing bias in brain morphometry analyses^54,55^. Low-quality scans were reacquired, and only the highest quality T1w/T2w pair per session was used. Additional 2-mm isotropic gradient echo and spin echo images were acquired for pseudo-transmit field computation^56^. For detailed protocol information, see Harms et al., 2018^57^.

#### Macaques

High-resolution 3D T1-weighted images were acquired on a GE MR750 3.0T scanner (General Electric Medical, Milwaukee WI) with an 8-channel brain array coil using an axial Inversion Recovery (IR) prepared fast gradient echo (fGRE) sequence (GE BRAVO) (0.55 × 0.55 × 0.8 mm, TI = 450 ms, TR = 8.684 ms, TE = 3.652 ms, FOV = 140 × 140 mm, flip angle = 12°, matrix = 256 × 256, thickness = 0.8 mm, gap = −0.4 mm, 80 percent field-of-view in phase encoding direction, bandwidth = 31.25 kHz, 2 averages). Structural T2-weighted images were acquired using a sagittal 3D CUBE FSE sequence (0.6 mm isotropic voxels, TR = 2500 ms, TE = 87 ms, FOV = 154 × 154 mm, flip angle = 90°, matrix = 256 × 256, 90 percent field of view in the phase encoding direction, slice thickness = 0.6 mm, gap = 0 mm, bandwidth = 62.5 kHz, ARC parallel imaging with a factor of 2 acceleration in both phase encoding and slice encoding directions). Animals were scanned under anesthesia. Subjects younger than 6 months were immobilized using inhalant isoflurane, while older subjects received ketamine hydrochloride (10 mg/kg I.M.) and dexdomitor (0.01 mg/kg I.M.). Heart rate and oxygen saturation were monitored throughout the experiment.

### Image Preprocessing

#### Humans

Preprocessed structural MRI data were provided as part of the Lifespan HCP Release 2.0 through the NDA (https://nda.nih.gov/). The data were processed using HCP Pipelines (version 4.0.1) within the QuNex container environment (qunex.yale.edu). As part of this pipeline, T1w and T2w volumes were processed through the *PreFreeSurfer* pipeline, which included gradient nonlinearity distortion correction, initial brain-extraction, and rigid registration into an anterior/posterior-commissure aligned “native” space. Each subject’s T2w volume was registered to their T1w volume using boundary-based registration (BBR)^58^, and the receiver coil bias field was corrected based on the smoothed square root of the product of the T1w and T2w images. Images were then registered to MNI template space^59^. The *FreeSurfer* pipeline (v6.0.0)^60^, optimized for use with high-spatial resolution (>1 mm isotropic) images, was used to compute the “white” and “pial” surfaces, including use of the T2w volume to optimize the pial surface placement. The values for the corpus callosum and ventricles were extracted from individual T1w and T2w images and calibrated using the group-averaged values of all subjects in the HCP dataset^61^.

#### Macaques

Preprocessed structural MRI data were provided as part of the UNC-Wisconsin Rhesus Macaque Neurodevelopment Database^47^. AutoSeg^62^ was used to perform the processing of the structural images. Bias field corrections were applied using the N4^63^ method for both T1 and T2 images^63^. T1 images were aligned to an external T1 atlas (Emory-UNC atlas at 12 months) using BRAINSfit’s rigid body, normalized mutual information registration. T2 images were aligned to the atlas-registered T1 images. These structural images were re-sampled in atlas space to 0.2375 × 0.2375 × 0.2375 mm resolution. Tissue segmentation was performed using the Atlas Based Classification4 (ABC) for white matter, gray matter, cerebrospinal fluid, and background^64^. Tissue segmentation results were used to generate binary brain masks that were corrected manually by an expert. Both T1w and T2w images were then warped to the NMT template space^27,65^. As was done for the human preprocessing, the values for the pons and ventricles were extracted from individual T1w and T2w images and calibrated using the group-averaged values of all subjects^61^.

and calibrated individually using the pons and ventricles.

### T1w/T2w ratio calculation

T1w/T2w maps were generated by dividing the T1w image by T2w image in standard template space (MNI for humans; NMT for macaques). This division cancels the signal intensity bias related to the sensitivity profile of the radio frequency receiver coils. The T1w/T2w ratio also enhances contrast associated with myelin content, as both images exhibit myelin-related contrast^25^ that is inverted in the T2w image relative to the T1w image. Individual T1w/T2w maps were parcellated using the Schaefer 400 parcellation^29^ for humans and the CHARM level6 139 parcellation for macaques^27,28^ to calculate the average T1w/T2w value for all vertices within each parcel. These T1w/T2w units are arbitrary, representing relative measures of intracortical myelin content that depend on the scanner field strength and sequence parameters, and should therefore not be directly compared across studies without appropriate normalization.

### Layer segmentation

The cortical gray matter was segmented using the LN2_LAYERS functions from LayNii^33^. The gray matter mask in template space (MNI for humans; NMT for macaques) was segmented according to the equi-volume principle^66^, which considers that outer layers have thinner volumes and produces depth bins of equal volume^33,66–68^. For Figures 1 and S2, the cortical gray matter was segmented into six equi-volume bins to maintain correspondence with the biological six layers. For Figures 2 and 3, it was segmented into seven bins, with the first and last bins removed to avoid contamination from white matter or external brain structures, resulting in five bins.

### Geodesic distance from default mode network regions

#### Humans

The geodesic distance along the cortical surface was calculated using tvb-gdist^69^ module (https://github.com/the-virtual-brain/tvb-gdist) that approximates the shortest path between two nodes on a triangular surface mesh. We selected parcels in the Schaefer 400 parcellation^29^ corresponding to the Default Mode Network (DMN) B from Yeo’s 17 networks^70^. Cortical nodes within these DMN parcels were clustered using k-means clustering, with 10% of the clusters randomly selected as seed nodes for efficiency. Each cortical node was then assigned a distance value based on the minimum geodesic distance along the “midthickness” surface to any of the seed nodes. Finally, the average of the minimum geodesic distances for nodes within each cortical parcel of the Schaefer 400 parcellation was calculated.

#### Macaques

The seed nodes were selected from five peak nodes from the principal gradient^31^, derived from axonal tract tracing decomposition, located within clusters corresponding to independent regions of the default mode network^31^. The same method for calculating geodesic distance in human data was applied. The average of the minimum geodesic distances for nodes within each cortical parcel of the CHARM level 6 parcellation ^27,28^ was calculated.

### Alignment with the Sensorimotor–Association (S-A) axis

#### Humans

We used the S-A axis derived by Sydnor and colleagues^30^. This map integrates various cortical hierarchies, including functional connectivity gradients, evolutionary cortical expansion patterns, anatomical ratios, allometric scaling, brain metabolism measures, perfusion indices, gene expression patterns, primary modes of brain function, cytoarchitectural similarity gradients, and cortical thickness.

#### Macaques

We used the S-A axis derived by Margulies and colleagues^31^. This map represents the principal gradient revealed through the decomposition of the axonal tract-tracing connectivity database CoCoMac^71,72^. To calculate the correlation between geodesic distance and the S-A axis in macaques, we aligned the CHARM level 6 parcellation^27,28^ with the Bonin-Bailey parcellation^73,74^ on Yerkes19 template^75^. We identified the most overlapping parcel in the Bonin-Bailey parcellation for each CHARM parcel. Parcels in the CHARM parcellation that did not overlap with any parcels in the Bonin-Bailey parcellation were left unassigned.

### Generalized Additive Models (GAMs)

To model both linear and non-linear relationships between the T1w/T2w ratio and age, we employed Generalized Additive Models (GAMs) implemented with the mgcv package in R^76^. In these models, the region-averaged T1w/T2w ratio served as the dependent variable, with age modeled as a smooth term, and gender included as linear covariates. Models were fitted separately for each cortical parcel, using thin plate regression splines as the basis set for the smooth term and the restricted maximum likelihood approach for selecting the smoothing parameter. The smooth term for age generated a spline representing each region’s developmental trajectory, with a maximum basis complexity (*k*) set to 3 to avoid overfitting.

For each regional GAM, we tested the significance of the association between the T1w/T2w ratio and age using an analysis of variance (ANOVA), comparing the full GAM model to a nested model without the age term. A significant result, as evaluated by the chi-squared test statistic, indicated that the inclusion of the age smooth term significantly reduced the residual deviance. Using the gratia package in R, we identified the specific age range(s) for each regional GAM where the T1w/T2w ratio showed significant changes by examining the first derivative of the age smooth function (Δ T1w/T2w ratio/Δ age) and determining when its simultaneous 95% confidence interval did not include zero (two-sided).

To quantify the magnitude and direction of the association between the T1w/T2w ratio and age, referred to as a region’s overall age effect, we measured effect size by calculating the partial *R^2^*comparing the full GAM model with a reduced model that excluded the age term. We signed the partial *R^2^* based on the average first derivative of the smooth function to indicate the effect’s direction. We sorted the parcels based on their S-A rank and divided them into three groups: sensorimotor, middle and association regions, each consisting of 133 parcels. We then performed the same GAM analysis on the average values across parcels within each group. For the macaque study, given the longitudinal design with multiple measurements obtained from the same monkeys over time, we accounted for within-subject correlation by including subject-specific random effects. This approach controls for variability arising from repeated measurements within the same individuals.

## Supplementary Results & Figures

### Validation of cortical myelin imaging with MBP expression in macaques

Although imaging modalities such as T1w/T2w ratio, magnetization transfer (MT), and myelin water fraction (MWF) can indirectly reflect myelin content across the cortical surface^24–26,32,42–44,77,78^, this relationship is imperfect^79,80^, and their specificity across cortical depths remains less established^18,81^. To assess the correlation between the T1w/T2w ratio and myelin content across both regions and cortical depth^10,39^, we compared it with the expression of myelin basic protein (MBP) in macaques using single-cell sequence^45^. Spatially, MBP expression was higher in sensorimotor regions than in association regions (Fig. S2A, B *R^2^* = 0.250, *F*(1, 131) = 24.350, *P* < 0.001). The T1w/T2w ratio showed a significant positive correlation with MBP expression across cortex (Fig. S2C *R^2^* = 0.216, F(1, 131) = 37.3, *P* < 0.001), consistent with prior findings ^24,42–44,77,78,82^.

For depth-wise validation, MBP expression was also significantly higher in deeper layers than in superficial layers (Fig. S2D, E, all ANOVAs *F* > 9.236, *P* < 0.001). To assess finer grained correspondence within individual cortical areas, we calculated the correlation coefficient between MBP and the T1w/T2w ratio for each CHARM parcel (Fig. S2F). Among the 86 parcels that had six histological layers identified in the MBP data, 88% (76 parcels) showed a significant correlation (*r* > 0.815, *P* < 0.050). A region-wise permutation test confirmed that this depth-dependent correlation is significantly higher than expected by random chance (*T* = −108.309, *P* < 0.001). These consistent depth-dependent patterns across broad cortical regions and the high proportion of significant correlations at the individual areas level validate the T1w/T2w ratio as a reliable marker for myelin content across both species and provide a robust method for assessing myelination variations across regions and depths.

**Supplementary Figure 1.**
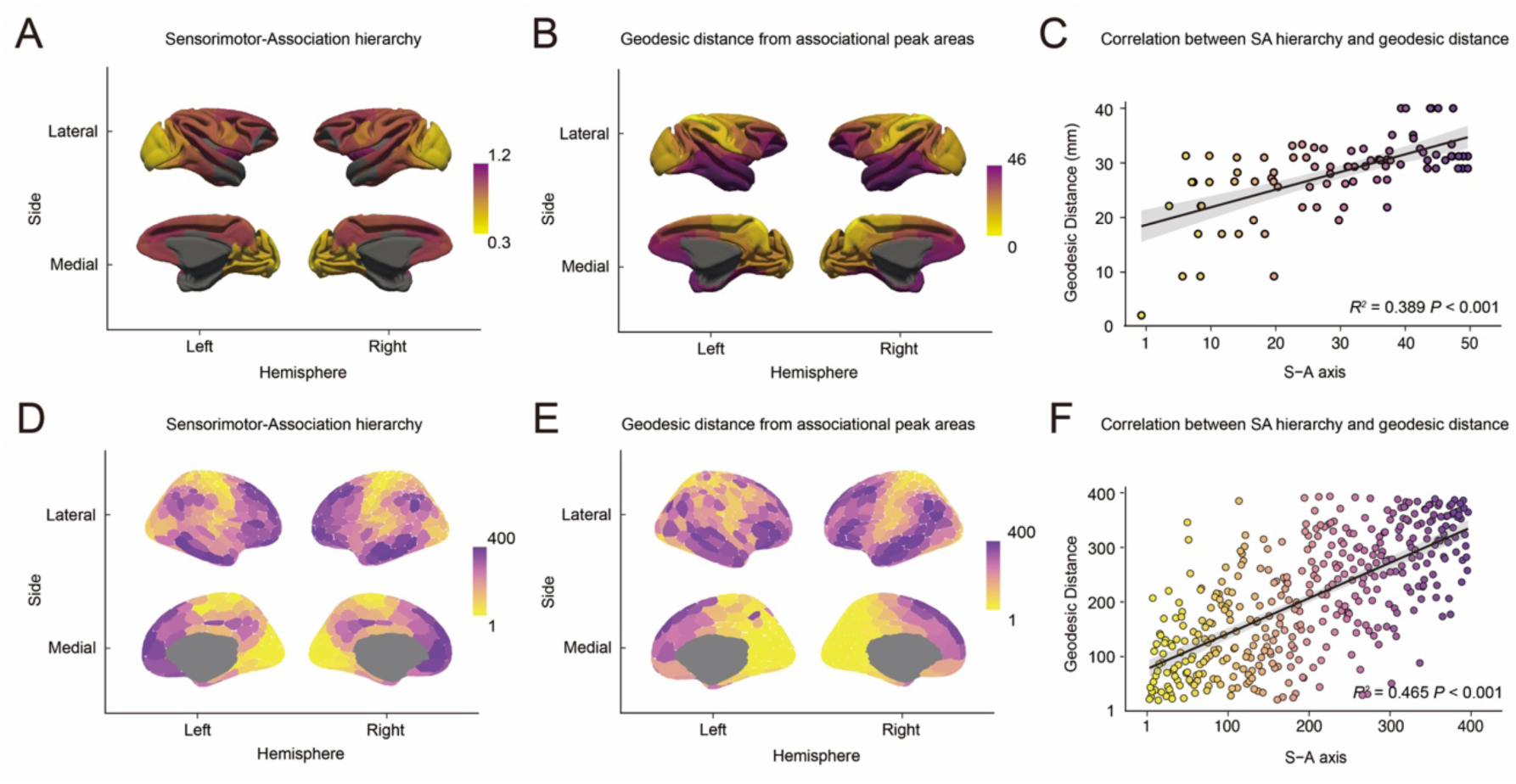
Correlation between geodesic distance and the sensorimotor-association axis. **(A, D)** Sensorimotor-association axis in macaques (A) and humans (D). **(B, E)** Geodesic distance from association regions in macaques (B) and humans (E). **(C, F)** Correlation between S-A axis and geodesic distance in macaques (C, *R^2^*= 0.389, *P* < 0.001) and humans (F, *R^2^* = 0.465, *P* < 0.001).

**Supplementary Figure 2.**
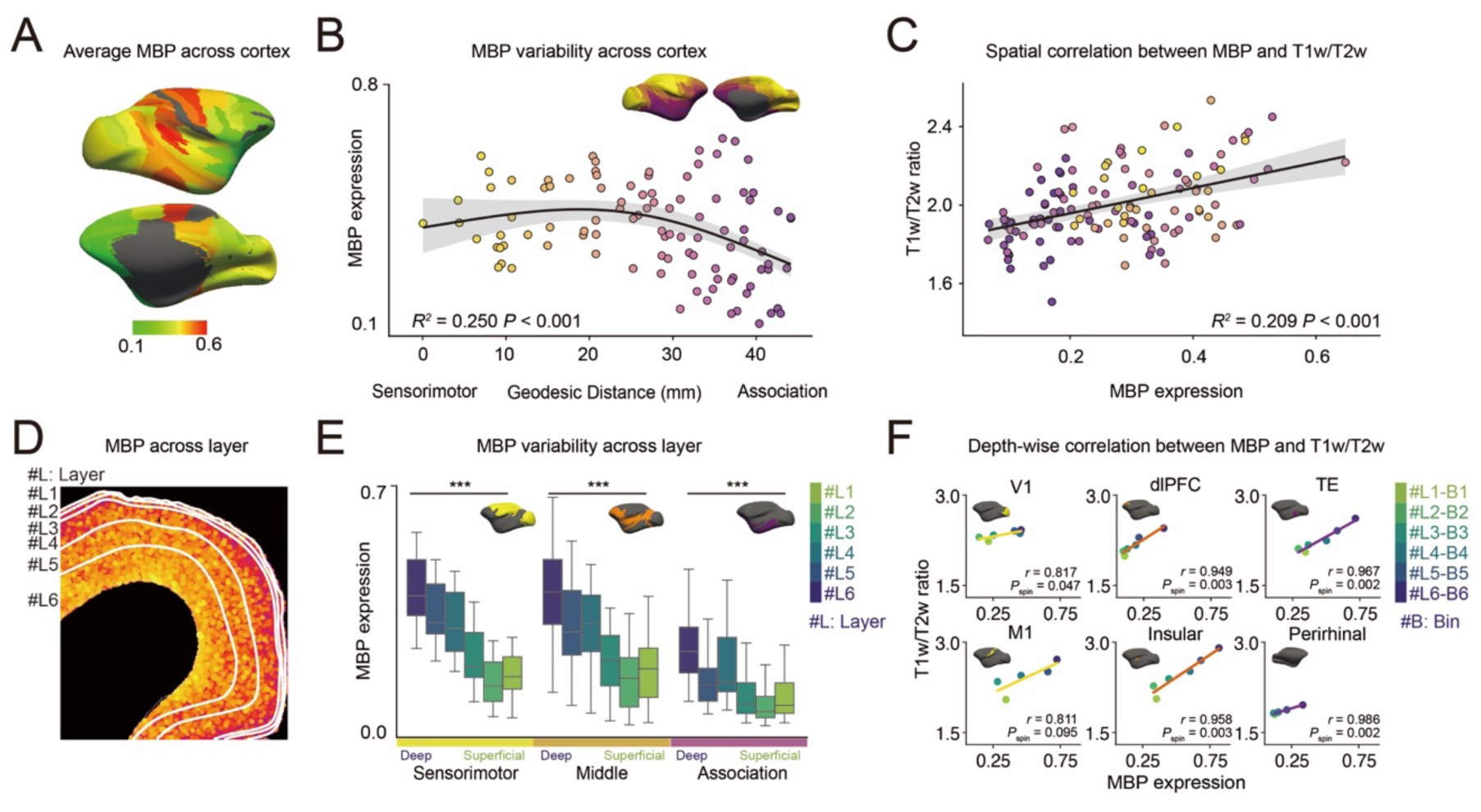
Correlation between MBP and T1w/T2w ratio across layers in macaques. **(A)** MBP expression across the cortex. **(B)** MBP expression along the geodesic distance from association regions. Brain regions are arranged on a spectrum from farther (yellow) to closer (purple) to the association centers. The GAM-predicted fit is displayed with a 95% confidence interval (*R^2^* = 0.250, *P* < 0.001). **(C)** Correlation between MBP expression and the T1w/T2w ratio across the cortex (*R^2^* = 0.209, *P* < 0.001). The regression line is displayed with a 95% confidence interval (*R^2^* = 0.209, *P* < 0.001). **(D)** Schematic illustration of MBP expression across cortical layers. **(E)** MBP expression in each layer for sensorimotor (yellow), middle (orange), and association (purple) regions. ANOVA \*\*\**P* < 0.001. **(F)** Correlation between MBP expression and T1w/T2w ratio across layers or bins in representative brain regions.

**Supplementary Figure 3.**
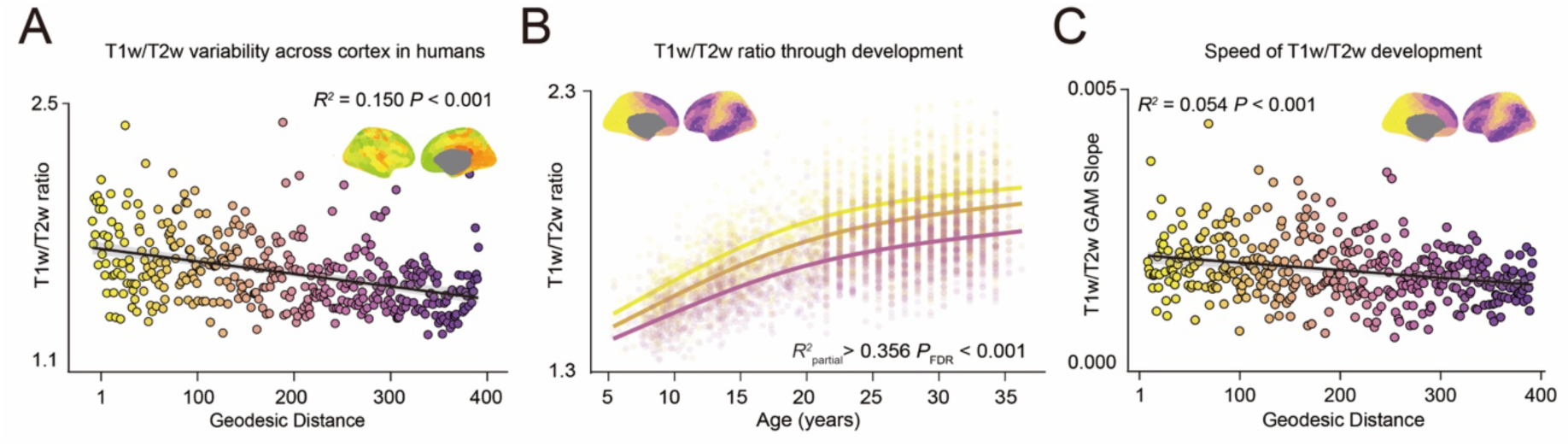
Development of T1w/T2w ratio along geodesic distance in humans. **(A)** T1w/T2w ratio across the cortex in humans. Brain parcels are aligned along geodesic distance from default mode network regions (*R²* = 0.150, *P* < 0.001). (B) Developmental trajectories of the T1w/T2w ratio for sensorimotor (yellow), middle (orange) and association (purple) cortical areas. Solid lines illustrate Generalized Additive Model (GAM)-predicted fits along with their 95% confidence intervals (Sensorimotor; *R*^2^_partial_ = 0.466, *P*_FDR_ < 0.001, Middle; *R^2^* _partial_ = 0.430, *P*_FDR_ < 0.001, Association; *R^2^* _partial_ = 0.356, *P*_FDR_ < 0.001). **(C)** GAM slope along the geodesic distance for humans. The GAM-predicted fit is displayed with a 95% confidence interval (*R²* = 0.054, *P* < 0.001).

## Notes

### Competing Interest Statement

The authors have declared no competing interest.

